# Antitumor Effects Of Phytochemicals Of Hot Three Plants On Gastric Adenocarcinoma

**DOI:** 10.1101/2021.11.21.469373

**Authors:** Narankhuu Ragchaasuren, Munkhtuya Tumurkhuu, Tserentsoo Byambaa, Uranbileg Bold, Enkhmaa Dagvadorj, Odontuya Gendaram, Tserendagva Dalkh

**Affiliations:** Mongolian Department of Medicine, Mongolian International University of Traditional Medicine, Mongolian National University of Medical Sciences, Ulaanbaatar, Mongolia; Department of Chemistry, School of Science, North Carolina State University, USA; Department of Molecular Biology and Genetics, School of Bio-Medicine, Mongolian National University of Medical Sciences, Ulaanbaatar, Mongolia; Department of Chemistry, Academy of Sciences, Ulaanbaatar, Mongolia

**Keywords:** Ranunculus, Ranunculaceae, thymoquinone, cancer, malignancy, tumor

## Abstract

**BACKGROUND:** Thymoquinone (TQ) and beta-sitosterol (BS), the bioactive constituents derived from the medicinal plant *Atragene sibirica* L, *Ranunculus repens* L, *Pulsatilla Bungeana* have been used for anti-cancer treatment in Traditional Mongolian Medicine. Recent studies reported that TQ, BS exhibited anti-proliferative effects on several cancer cell lines. This study was performed to investigate the antitumor effects of phytochemicals on gastric adenocarcinoma *in vitro*.

**METHODS:** To this aim, we performed a cell apoptosis assay WST1 and several cancer-related gene expressions (*Bcl, BIRC5, p53, BAX*). NCI-N87, MKN74 cells were seeded at a density of 5×10^4^ cells per well in 96-well culture plates. After overnight incubation, the fresh medium containing different concentrations of TQ (20, 40, 60, 80, and 100 mM/ml) and beta-sitosterol (10, 1, 0.1, 0.001, 0.001 mM/ml) were applied. Following a 24-h incubation, the cell viability was determined by WST-1 assay. We also quantified the expression levels of mRNAs of these genes through qRT-PCR experiments.

**RESULTS:** Our results showed that TQ induced a higher percentage of cell apoptosis in the human gastric adenocarcinoma cell lines compared to that of control. We observed a significant 4.8-fold change in *BAX/BCL2* ratio. These increases were highly correlated with a concentration-dependent manner (p=0.00178). Moreover, the expression of *p53* and *BIRC5* was downregulated in MKN74 cells after treatment with TQ.

**CONCLUSIONS:** Our results demonstrate that TQ effectively inhibits cell proliferation through several gene expressions in vitro. Moreover, inhibition of the downstream molecule of these genes would explain the underlying mechanism of the antitumor activity in cancer cell lines.

## INTRODUCTION

Gastric cancer (GC) is one of the most prevalent cancer in the world, accounting for the fourth case for cancer-related mortality, where the median survival rate is less than 12 months for the advanced stage^1,2^. GC is two to three times more prevalent among men ^3^ and the rate of GC is very high in Asian countries^4^. GC develops from precancerous lesions, chronic gastritis as well as helicobacter pylori-induced inflammation ^5^. Surgery is currently considered to be the only treatment option, with the radio and chemotherapy the treatment success reaches more than 95%. However, the low rate of early diagnosis closes the chances of the treatment, reaching only a 40% survival rate ^6^. Therefore, the development of novel drugs against gastric cancer and early detection methods are in critical need for better diagnosis and treatment of the patients.

With the advanced genetic tools, cancer has been studied at the genetic level ^7^. Differentially expressed genes (DEG) are studied well in disease prognosis and drug screening process ^8^. DNA microarray technology has played an important role in identifying genetic causes for GC cases ^9^, but the underlying mechanisms are still not clear.

Previous studies revealed that some DEGs are responsible for carcinoid lesions in the stomach, which are induced by elevations of gene expression such as *BCL-2* ^10^, *BIRC5* (survivin) ^11,12^, *Bax* ^9^, *p53*^13^, *ABCB1*^14^*, UHRF1*^15^, and *DNMT1*^16^. These genes are involved by the Hippo BIRC5 pathway, p53 signaling pathway, intrinsic apoptotic pathway that interacts with XIAP and DIABLO leading to caspase-3 and caspase-9 inactivation, or regulators of apoptosis, WNT pathway, UHRF1-dependent regulation, and DNA methyl-transferase pathway.

The gastric cancer developmental process is influenced by both genetic and environmental factors. Approximately half of the cancer incidents might be provoked by environmental agents, mostly dietary habits, bacterial infection and, social behavior. The development and progression of malignant tumors is a multiannual and multistage process. Especially gastric cancer, a multifactorial disease influenced by genetic and environmental factors is usually diagnosed in the advanced stage of the disease ^17,18^.

In our present study, we have investigated three medicinal plants used in Mongolian Traditional Medicine against gastric cancer, named *Atragene sibirica* L, *Ranunculus repens* L, *Pulsatilla Bungeana* L. Bio-active products from these plants were extracted using vacuum extract methods and dissolved in alcohol, organic, and water solutions. Bio-materials dissolved in inorganic and organic solvents were used to screen cell toxicity depending on the different concentrations. For this purpose, we used two well-differentiated cell lines, MKN74, NCI-N87 cells, and several gene expressions were screened for cancer-related activities. Also, we chose two biologically active ingredients, thymoquinone, and beta-sitosterol.

Thymoquinone (2-methyl-5-isopropyl-1,4-benzoquinone) is a phytochemical product, has been used in different traditional medical applications found in Chinese, Mongolian and Ayurvedic Medicine found in Arabic, Mediterranean, African countries. Thymoquinone has been shown to exert anti-inflammatory, antidiabetic, hepatoprotective, renal-protective, and stomach-protective as well as anti-cancerous effect ^19^. TQ inhibited colony formation and cell migration ability of the MGC80-3 and SGC-7901 cells gastric cancer cells and down-regulated the expression of the mesenchymal genes such as *N-cadherin, Vimentin*, and TWIST ^20^.

Beta-sitosterol is an essential phytochemical that is enriched in plant food. The most common plant sterols in foods are beta-sitosterol, campesterol, and stigmasterol, representing about 50-65%, 10-40%, and 0-35% of the total phytosterols, respectively. It has been shown to exert antiproliferative effects on the different types of cancers of the colon, breast, and prostate, but its effect on gastric adenocarcinoma cells *in vitro* is unknown^20^. These phytochemicals’ effects on cancer-related gene expressions are not well studied as they could serve as vital sources for novel drugs for cancer therapy.

## MATERIAL METHODS

*Atragene sibirica* L, *Ranunculus repens* L, *Pulsatilla Bungeana* L. plants were harvested from the southern part of Mongolia from August 2020 to August 2021. These plants were dried and biologically active materials were dissolved in different inorganic and organic solvents. These vials were shipped to IUHW, Chiba, Japan for cell viability and gene expression assays.

### CELL CULTURE

Human MKN74 and NCI-N87 are well-differentiated adenocarcinoma human gastric cancer cell lines that were generously provided by Prof. Hiroshi Tazawa (Okayama University, Japan). MKN74 and NCI-N87 cells were maintained in modified Eagle’s medium containing 10% fetal bovine serum (FBS), 1% penicillin-streptomycin, and 1% glutamine at 37°C with 5% CO2. MKN74 and NCI-N87 cells were seeded at a density of 5×10^4^ cells per well in 96-well culture plates. After overnight incubation, the medium was removed and replaced with a fresh medium containing different concentrations of thymoquinone (20, 40, 60, 80, and 100 μM/ml). Also, the serum-containing medium was replaced by a fresh medium containing 0.001, 0.01, 0.1, 1, and 10 μM/ml β-sitosterol dissolved in DMSO. The final working concentration of DMSO was <0.1%. Also, 96% of ethanol, n-Hexane, Chloroform (pH=8), 80% ethanol, Dichloromethane, ethyl acetate, butanol, water solutions of three plants were screened in 25, 50 μg/μl concentrations for cell viability assay.

### CELL VIABILITY ASSAY

Following a 24-h incubation with these products, the cell viability was determined by WST-1 assay (CELL PRO-RO Roche, Japan) according to the manufacturer’s instructions. Briefly, WST1 solution containing tetrazolium salt was added to cells in 96-well plates, the cells were then incubated at 37°C for 4 hours, and absorbance was measured at 420-480 nm using an MRX Revelation 96-well multi scanner (Dynex Technologies, Chantilly, VA, USA). This experiment was repeated three times.

### GENE EXPRESSIONS ASSAY

Thymoquinone and beta-sitosterol were screened for gene expression assay. The MKN74 and NCI-N87 cells were seeded at a density of 5×10^4^ cells per well in 96-well culture plates. After overnight incubation, the medium was removed and replaced with a fresh medium containing thymoquinone and β-sitosterol dissolved in DMSO for 24 h. For preparation, 10uL β-mercaptoethanol to 1ml RLT, 4 volumes of 100% ethanol to RPE were added in advance. Cells were washed with 1XPBS three times, 600 uL of Buffer RLT were added and harvested cells by scraping. The cells were shredded by QIAshedder homogenizer (Qiagen USA, cat No.74054), centrifuged with 15000rpm for 2 min at rt. RNeasy Minikit (Qiagen USA, cat. No. 74034) was used according to the manufacturer’s protocol. Extracted RNA concentration was checked by Nanodrop 2000 spectrophotometer. Cellular RNA was transferred into cDNA using a High-Capacity cDNA Reverse Transcription Kit with RNase Inhibitor (Applied Biosystems, cat No. 374966). Thermal cycling was conducted in 25°C 10min, 37°C 120min, 85°C 5min and transferred to 4°C. cDNA was stored at −20°C for further experiment. qRT-PCR was done with Brilliant III Ultra-Fast SYBR Green (Agilent) with *ACTB* gene as homeobox gene. We used *BCL-2* ^10^, *BIRC5* (survivin) ^11,12^, *Bax* ^9^, *p53*^13^, *ABCB1*^14^*, UHRF1*^15^, and *DNMT1*^16^ genes for apoptotic effects after thymoquinone and beta-sitosterol treatment. Gene-specific primers were designed by Primer3 software (https://bioinfo.ut.ee/primer3-0.4.0/primer3/input.htm). Two times Brilliant III Ultra-Fast SYBR Green QPCR Master Mix with Low, Primer 1,2 (10 pmol/l, final 500 nM), RNase free water, cDNA (10ng/μL) with the total volume of 20 was used for the RT-PCR reaction. cDNA concentration was adjusted to 10ng/μL, by adding 80μL TE to the solution of 1μg/20μL (1μg/100μL = 10ng/μL). Expressions of the cDNA were calculated by the delta-delta Ct method (ΔΔCT). TaqMan Fast Advanced Master Mix, 1mL (ABI-Thermo Fisher 4444557), Tris-EDTA solution 100x (Sigma T9285-100ML), MicroAmp Optical Adhesive Film (25) (Life Technologies 4360954) were used.

## RESULTS

Our results showed that *Ranunculus repens* L and *Atragene sibirica* L increases cellular apoptosis depending on solvent as well as concentration. The cancer cell lines were significantly decreased (p>0.0002) after the treatment with alcoholic (96% and 80% ethanol) and organic solvents as well as water solutions (Fig 1.) This phenomenon was also observed in *Atragene sibirica* L samples (Fig 2.) It showed extracts of *Atragene sibirica* L with ethanol solvents had apoptotic effects on the gastric cancer cell lines but not the other solvents. We screened for gene expressions of *BCL-2*, *BIRC5*, *Bax*, *p53*, *ABCB1, UHRF1, DNMT1,* and *HDAC* after the treatment of beta-sitosterol and thymoquinone with concentrations of 0.001, 0.01, 0.1, 1, 10 μM/ml and 20, 40, 60, 80, 100 μM/ml for 24 h (Fig 3.). Genes of *BCL-2*, *BIRC5*, *Bax*, and *p53* were highly activated after the treatment of thymoquinone in concentrations of 80, 100mM/mL (p>0.05), but not in lower concentrations of 20, 40, and 60 mM/mL (Fig 4.), and also this phenomenon was not observed in beta-sitosterol samples in all concentrations (Fig 5.) Genes expressions were normalized with Homeobox gene *ACTB* and delta-delta CT data was used for comparison. TQ induced a higher percentage of cell apoptosis in the human gastric adenocarcinoma cell lines compared to that of control. We observed a significant 4.8-fold change in *BAX/BCL2* ratio. These increases were highly correlated with a concentration-dependent manner (p=0.00178). Moreover, the expression of *p53* and *BIRC5* was downregulated in MKN74 cells after treatment with TQ.

**Fig 1.**
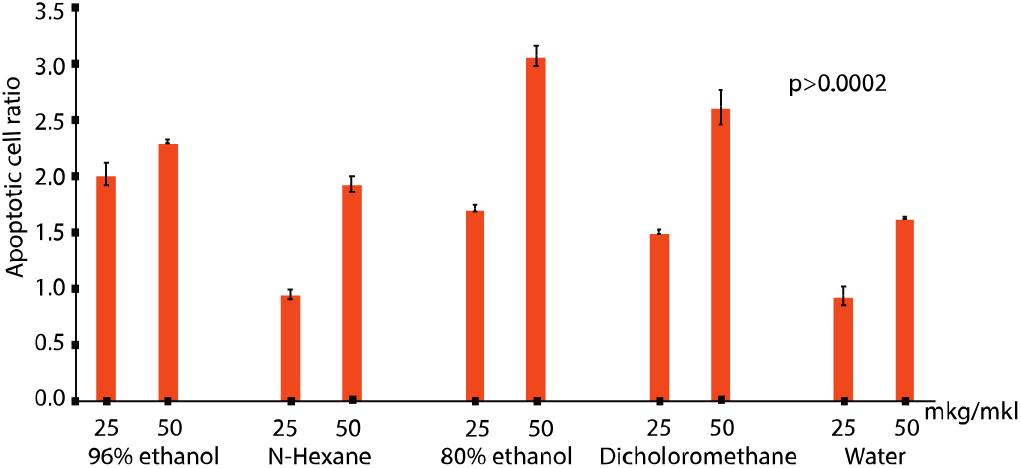
Cell apoptotic ratio after the treatment of *Ranunculus repens L* solutions.

**Fig 2.**
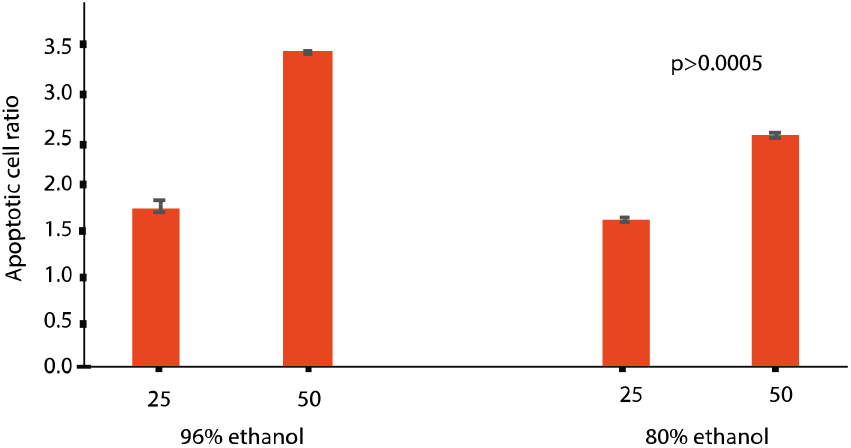
Cell apoptotic ratio after the treatment of *Atragene sibirica* L solutions.

**Fig 3.**
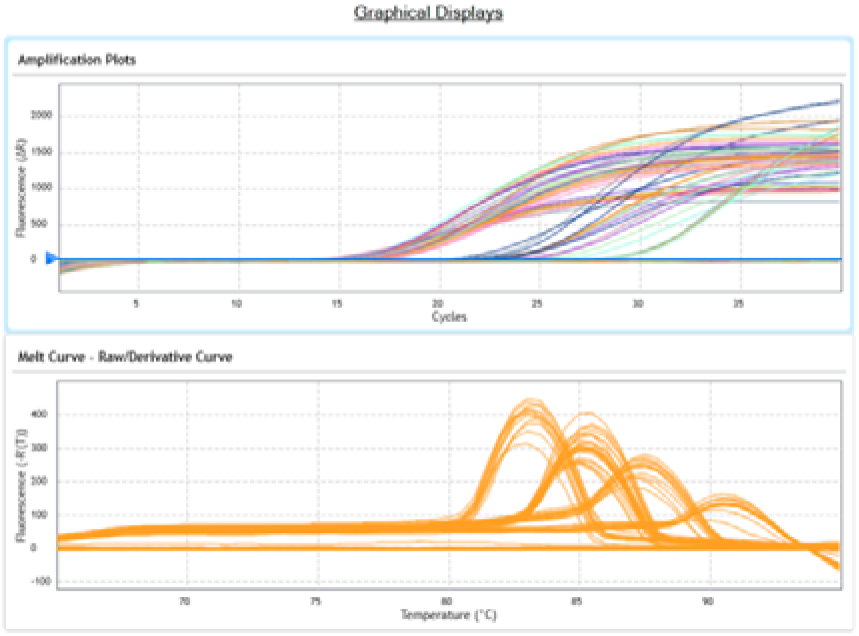
RT-PCR graphical results of the genes

**Fig 4.**
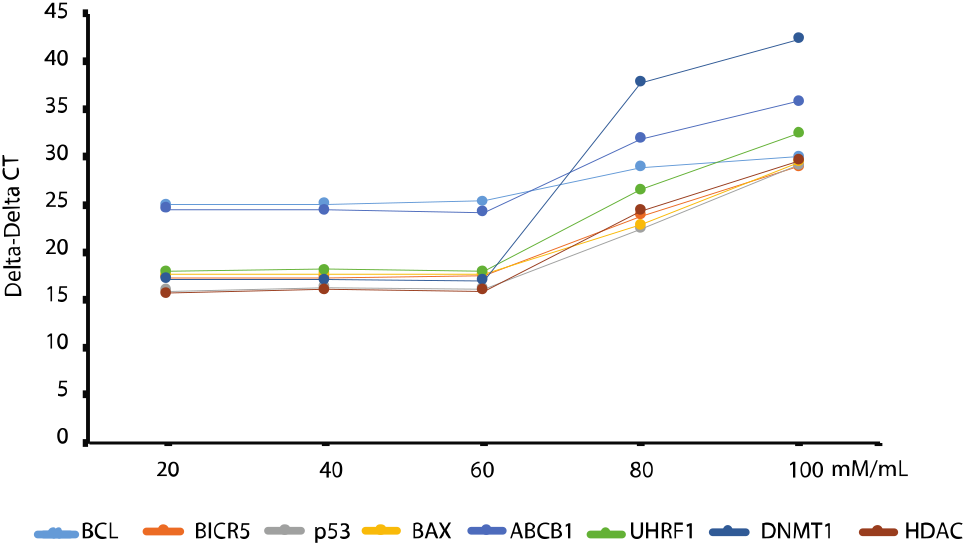
Genes expressions elevated after treatment of Thymoquinone in different concentrations.

**Fig 5.**
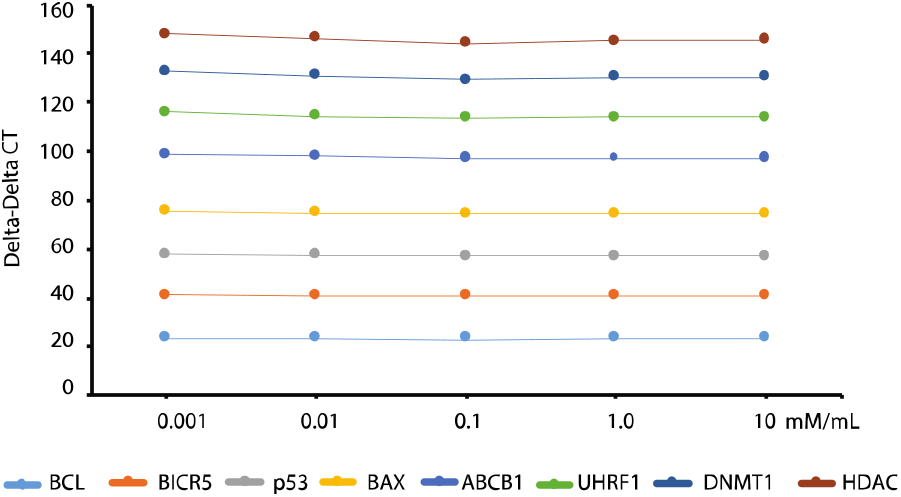
Genes expressions elevated after treatment of beta-sitosterol in different concentrations.

## DISCUSSION

Gastric cancer (GC) is one of the most common malignancies in humans and it is the fourth leading cause of cancer-related morbidity. As carcinogenesis is a multistage, multifactorial disease, where environmental and genetic factors play significant roles ^2^. About half of the GC cases might be provoked by environmental factors, dietary habits, and social behavior. It is different regarding sex and geography, where men have two to three times higher risk than women, and more than 70% of cases occur in developing countries ^3^. Countries with the highest probability for GC development encompass regions like Central and South America, Eastern Europe, and Asia. Mongolia has a high burden of non-communicable diseases, where cancer-related diseases account for the second biggest cause of morbidity ^21^. Mongolians have the highest age-standardized cancer-related deaths in the Asia Pacific region, which is explained by diagnosis at a later stage, common cancers associated with poor survival, and limited quality of health services ^21^. In a global cancer survival study covering 67 countries, Mongolia had among the lowest 5-year survival estimates for lung and liver cancers, about 7– 9% ^21^. Therefore, we should improve diagnostic and treatment options including alternative treatments for GC, using Traditional Mongolian Medicine (TMM). TMM has hundreds of years of development, but unfortunately, it was used in small quantities because of political reasons, which was abolished during 1921-1990. Lately, TMM is started greatly accepted among Mongolians because of its high benefits for non-communicable diseases. We chose “Hot three plants” which was prescribed for GC patients. These three plants are *Atragene sibirica* L, *Ranunculus repens* L, *Pulsatilla Bungeana* L, which have many different biological active products such as saponins, tannins, phenols, flavonoids, and alkaloids. Saponins are known for their anti-inflammatory, anti-oxidant, antirheumatic, and antimicrobial effects. Ranunculus family plants are known with rich amounts of thymoquinone (30% - 48%), thymohydroquinone, dithymoquinone, p-cymene (7% −15%), carvacrol (6% −12%), 4-terpineol (2% −7%), tanethol (1% −4%), longifolene (1% −8%), α-pinene and thymol ^22^. In this study, we chose a thymoquinone as a candidate treatment agency for GC, since it is known to harbor anti-inflammatory and anticancer effects ^23,24^. Thymoquinone effects were observed in our study depending on different concentrations. We observed a significant 4.8-fold change in *BAX/BCL2* ratio. These increases were highly correlated with a concentration-dependent manner (p=0.00178).

Beta-sitosterol is used in traditional medicine in Vietnam. The experimental results in this study revealed that β-sitosterol (β-S) and β-sitosterol-glucoside (β-SG) were the main ingredients of the *I. zollingeriana* extract ^25^. We used the pure extracts of Beta-sitosterol in our study to see the corresponding gene expression alterations in gastric cell lines and found that it increases apoptotic reactions based on concentrations but did not observe the chosen gene alterations. We explain it with different gene expressions that could be examined to see the apoptotic reactions after the treatment of beta-sitosterol. Also, it could be explained by a significant amount of beta-sitosterol or longer time treatment is necessary to provoke the treatment effect.

In our study, we screened *BCL-2*, *BIRC5*, *Bax*, *p53*, *ABCB1, UHRF1, DNMT1,* and *HDAC* gene activity after the TQ and BS treatment based on possible cell signal transduction pathway. *BIRC5* gene activity is regulated by the Hippo BIRC5 pathway, which is an anti-apoptotic downstream target gene of the Hippo pathway that plays a role in cell growth and apoptosis^25^. BCL-2 family proteins are the regulators of apoptosis, but also have other functions. This family of interacting partners includes inhibitors and inducers of cell death. Together they regulate and mediate the cell death process by which mitochondria contribute to cell death known as the intrinsic apoptosis pathway ^26^. BCL2 proteins cooperate with Bax proteins in apoptotic processes ^27^. *UHRF1, DNMT1,* and *HDAC* genes are also involved in the cell division, DNA methylation, posttranslational modifications, and intrinsic apoptotic pathway ^28–30^. These gene activities are altered in TQ and BS treated gastric adenocarcinoma cell lines, which proves our hypothesis to use it as an alternative therapeutic method for GC-related diseases by tea extracts. In the future, this research could be fundamental for making tea from these plants for everyday usage in Mongolia.

## CONCLUSION

1. Ethanol and N-Hexane extracts of *Ranunculus Repens L*, water extracts of P*ulsatilla Bungeana* L, ethanol extracts of *Atragene Sibirica* L contain biologically active ingredients that activate cancer cell apoptosis in a concentration-dependent manner.
2. TQ induced a higher percentage of cell apoptosis in the human gastric adenocarcinoma cell lines compared to that of control, where was observed a significant 4.8-fold change in *BAX/BCL2* ratio. These increases were highly correlated with a concentration-dependent manner (p=0.00178). Moreover, the expression of *p53* and *BIRC5* was downregulated in MKN74 cells after treatment with TQ.
3. Our results demonstrate that TQ effectively inhibits cell proliferation through several gene expressions in vitro. Moreover, inhibition of the downstream molecule of these genes would explain the underlying mechanism of the antitumor activity in cancer cell lines.

## ACKNOWLEDGEMENT

This publication resulted from research supported by the Mongolian Foundation for Science and Technology (2019-34) , Mongolian National University of Medical Sciences during 2019-2021. The content is solely the responsibility of the authors and does not necessarily represent the official views of the MFSC, Mongolia.

## REFERENCES

1. Zhang X, Zhang P. Gastric cancer: somatic genetics as a guide to therapy. J Med Genet. 2017;54(5):305–312. doi:10.1136/JMEDGENET-2016-104171

2. Machlowska J, Baj J, Sitarz M, Maciejewski R, Sitarz R. Gastric Cancer: Epidemiology, Risk Factors, Classification, Genomic Characteristics, and Treatment Strategies. Int J Mol Sci. 2020;21(11). doi:10.3390/IJMS21114012

3. Ferlay J, Shin H-R, Bray F, Forman D, Mathers C, Parkin DM. Estimates of worldwide burden of cancer in 2008: GLOBOCAN 2008. Int J Cancer. 2010;127(12):2893–2917. doi:10.1002/IJC.25516

4. Aiwen Wu JJ. The 7th National Gastric Cancer Academic Conference: focus on translational research in gastric cancer - Wu - Translational Gastrointestinal Cancer. / Vol 1, No 3 (October 2012). Accessed October 29, 2021. https://tgc.amegroups.com/article/view/1108/1456

5. Wang F, Meng W, Wang B, Qiao L. Helicobacter pylori-induced gastric inflammation and gastric cancer. Cancer Lett. 2014;345(2):196–202. doi:10.1016/J.CANLET.2013.08.016

6. Z S, YW, J Y, D Y, X F. Progress in the treatment of advanced gastric cancer. Tumour Biol. 2017;39(7). doi:10.1177/1010428317714626

7. Chong X, Peng R, Sun Y, Zhang L, Zhang Z. Identification of Key Genes in Gastric Cancer by Bioinformatics Analysis. Biomed Res Int. 2020;2020. doi:10.1155/2020/7658230

8. Sp P. From next-generation resequencing reads to a high-quality variant data set. Heredity (Edinb). 2017;118(2):111–124. doi:10.1038/HDY.2016.102

9. D’ Angelo G, Rienzo T Di, Ojetti V. Microarray analysis in gastric cancer: A review. http://www.wjgnet.com/. 2014;20(34):11972–11976. doi:10.3748/WJG.V20.I34.11972

10. C S, Y H, HL, et al. HOXA10 induces BCL2 expression, inhibits apoptosis, and promotes cell proliferation in gastric cancer. Cancer Med. 2019;8(12):5651–5661. doi:10.1002/CAM4.2440

11. Y Y, Z L, Y C, et al. Downregulation of TRIM27 suppresses gastric cancer cell proliferation via inhibition of the Hippo-BIRC5 pathway. Pathol Res Pract. 2020;216(9). doi:10.1016/J.PRP.2020.153048

12. J Z, X L, J Z, L W. Dysregulation of miR-195-5p/-218-5p/BIRC5 axis predicts a poor prognosis in patients with gastric cancer. J Biol Regul Homeost Agents. 2019;33(5):1377–1386. doi:10.23812/19-146-A

13. Gh W, X W. lncRNA MEG3 inhibitproliferation and metastasis of gastric cancer via p53 signaling pathway. Eur Rev Med Pharmacol Sci. 2017;21(17):3850–3856. Accessed October 29, 2021. https://pubmed.ncbi.nlm.nih.gov/28975980/

14. J de O, Av F, Ra N, Ct O, M de SS, Nm F. Association between ABCB1 immunohistochemical expression and overall survival in gastric cancer patients. Asian Pac J Cancer Prev. 2014;15(16):6935–6938. doi:10.7314/APJCP.2014.15.16.6935

15. H Z, YS, C Y, X W. UHRF1 mediates cell migration and invasion of gastric cancer. Biosci Rep. 2018;38(6). doi:10.1042/BSR20181065

16. HL, W L, S L, et al. DNMT1, DNMT3A and DNMT3B Polymorphisms Associated With Gastric Cancer Risk: A Systematic Review and Meta-analysis. EBioMedicine. 2016;13:125–131. doi:10.1016/J.EBIOM.2016.10.028

17. Ar Y, K BL, P B, M R, Z K. Risk Factors for Gastric Cancer: A Systematic Review. Asian Pac J Cancer Prev. 2018;19(3):591–603. doi:10.22034/APJCP.2018.19.3.591

18. Machlowska J, Baj J, Sitarz M, Maciejewski R, Sitarz R. Gastric Cancer: Epidemiology, Risk Factors, Classification, Genomic Characteristics, and Treatment Strategies. Int J Mol Sci. 2020;21(11). doi:10.3390/IJMS21114012

19. Khan MA, Tania M, Fu S, Fu J. Thymoquinone, as an anticancer molecule: from basic research to clinical investigation. Oncotarget. 2017;8(31):51907–51919. Accessed November 1, 2021. www.impactjournals.com/oncotarget/

20. Y Z, Sk C, G Q, T L, H C. Beta-sitosterol inhibits cell growth and induces apoptosis in SGC-7901 human stomach cancer cells. J Agric Food Chem. 2009;57(12):5211–5218. doi:10.1021/JF803878N

21. Chimed T, Sandagdorj T, Znaor A, et al. Cancer incidence and cancer control in Mongolia: Results from the National Cancer Registry 2008–12. Int J Cancer. 2017;140(2):302–309. doi:10.1002/IJC.30463

22. Begum S, Mannan A. A Review on Nigella sativa: A Marvel Herb. J Drug Deliv Ther. 2020;10(2). doi:10.22270/jddt.v10i2.3913

23. Imran M, Rauf A, Khan IA, et al. Thymoquinone: A novel strategy to combat cancer: A review. Biomed Pharmacother. 2018;106:390–402. doi:10.1016/J.BIOPHA.2018.06.159

24. Almatroodi SA, Almatroudi A, Alsahli MA, Khan AA, Rahmani AH. Thymoquinone, an Active Compound of Nigella sativa: Role in Prevention and Treatment of Cancer. Curr Pharm Biotechnol. 2020;21(11):1028–1041. doi:10.2174/1389201021666200416092743

25. Vo TK, Ta QTH, Chu QT, Nguyen TT, Vo VG. Anti-Hepatocellular-Cancer Activity Exerted by β-Sitosterol and β-Sitosterol-Glucoside from Indigofera zollingeriana Miq. Molecules. 2020;25(13). doi:10.3390/MOLECULES25133021

26. Marie Hardwick J, Soane L. Multiple Functions of BCL-2 Family Proteins. Cold Spring Harb Perspect Biol. 2013;5(2). doi:10.1101/CSHPERSPECT.A008722

27. Maes ME, Schlamp CL, Nickells RW. BAX to basics: How the BCL2 gene family controls the death of retinal ganglion cells. Prog Retin Eye Res. 2017;57:1. doi:10.1016/J.PRETEYERES.2017.01.002

28. Li Y, Seto E. HDACs and HDAC Inhibitors in Cancer Development and Therapy. Cold Spring Harb Perspect Med. 2016;6(10). doi:10.1101/CSHPERSPECT.A026831

29. Jorgensen BG, Berent RM, Ha SE, et al. DNA methylation, through DNMT1, has an essential role in the development of gastrointestinal smooth muscle cells and disease. Cell Death Dis. 2018;9(5). doi:10.1038/S41419-018-0495-Z

30. Patnaik D, Estève PO, Pradhan S. Targeting the SET and RING-associated (SRA) domain of ubiquitin-like, PD and ring finger–containing 1 (UHRF1) for anti-cancer drug development. Oncotarget. 2018;9(40):26243. doi:10.18632/ONCOTARGET.25425

